# Maximizing glycoproteomics results through an integrated PASEF workflow

**DOI:** 10.1101/2023.12.21.570555

**Authors:** Melissa Baerenfaenger, Merel A Post, Fokje Zijlstra, Alain J van Gool, Dirk J Lefeber, Hans JCT Wessels

**Author notes:** **Corresponding Author** Hans JCT Wessels - Translational Metabolic Laboratory, Department of Human Genetics, Radboud Institute for Molecular Life Sciences, Radboud University Medical Center, 6525 GA Nijmegen, The Netherlands. These authors contributed equally to this work and share first authorship (Melissa Baerenfaenger, Merel A Post).

## Abstract

Glycoproteins play important roles in numerous physiological processes and are often implicated in disease. Analysis of site-specific protein glycobiology through glycoproteomics is evolving rapidly in recent years thanks to hardware and software innovations. Particularly, the introduction of Parallel Accumulation Serial Fragmentation (PASEF) on hybrid trapped ion mobility time-of-flight mass spectrometry instruments combined deep proteome sequencing with separation of (near-)isobaric precursor ions or converging isotope envelopes through ion mobility separation. However, reported use of PASEF in integrated glycoproteomics workflows to comprehensively capture the glycoproteome is still limited. To this end, we developed an integrated methodology using the timsTOF Pro 2 to enhance N-glycopeptide identifications in complex mixtures. We systematically optimized the ion optics tuning, collision energies, mobility isolation width and the use of do-pant-enriched nitrogen gas (DEN). Thus, we obtained a marked increase in unique glycopeptide identification rates compared to standard proteomics settings showcasing our results on a large set of glycopeptides. With short liquid chromatography gradients of 30 minutes, we increased the number of unique N-glycopeptide identifications in human plasma samples from around 100 identifications under standard proteomics condition to up to 1500 with our optimized glycoproteomics approach, highlighting the need for tailored optimizations to obtain comprehensive data.

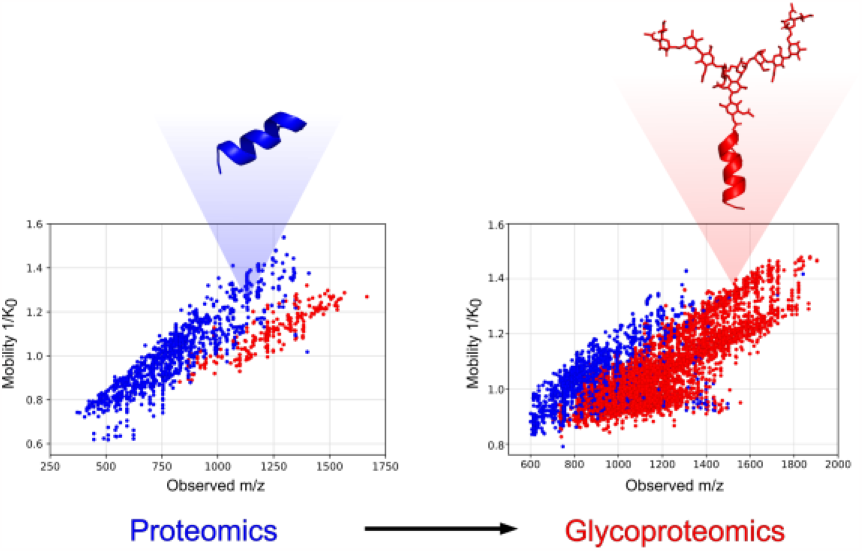

## INTRODUCTION

Glycosylation is the most common post-translational modification of human proteins.^1-2^ Its implication in all crucial processes of cellular life has resulted in a growing interest to analyze glycosylation to answer biological and clinical questions.^1, 3-6^ However, studying protein glycosylation is challenging due to the inherent structural complexity caused by micro- and macroheterogeneity and the existence of various glycan isomers.^7-9^ As a result, the field of glycoproteomics is lacking behind its bottom-up proteomics counterpart where mass spectrometry (MS) has proven to be a powerful tool for identifying many thousands of proteins in a single LC-MS/MS experiment.^10-12^ These advancements were supported by well-established workflows for sample preparation, MS acquisition and data interpretation for bottom-up proteomics.^13-14^ In comparison, the holistic analysis of glycopeptides remains challenging due to physicochemical properties of glycopeptides and immature technology that lacks standardized workflows. Hence, MS identification rates of glycopeptides from complex mixtures are much lower compared to peptide identifications in bottom-up experiments.^15-16^ When compared to bottom-up proteomics, glycoproteomics approaches require a tailored approach that favors the detection of glycopeptides over peptides. After tryptic digestion of proteins and glycoproteins, the large quantity of peptides in the sample mixture can suppress the detection of less abundant glycopeptides. Therefore, a glycopeptide enrichment can be performed to increase the number of glycopeptides and deplete non-glycosylated peptides.^17-18^

However, further challenges arise in the mass spectrometric detection of glycopeptides. For example, the hydrophilicity of glycopeptides decreases their ionization efficiency in positive ionization mode.^19^ To increase ionization efficiency, especially in samples with a large amount of non-glycosylated peptides that could suppress signals of glycopeptides, several strategies have been developed. The use of dopant-enriched nitrogen gas (DEN) has shown to significantly increase desolvation and ionization of hydrophilic analytes such as glycans, glycopeptides or glycoproteins.^20-23^ Here, an organic modifier, such as acetonitrile (ACN) or primary alcohols are used to enrich the nebulizer gas during electrospray ionization which is not only favorable for the signal intensity of glycopeptides but can also influence their charge state distributions. The use of acetonitrile as DEN has shown to increase the charge of glycopeptides from 2+ to 3+ toward 3+ to 4+ or higher in combination with reversed phase nanoLC-MS.^20^ This is a significant advantage as higher charge states of glycopeptides also address another obstacle in the MS/MS detection of glycopeptides. Similar to peptides, higher charge states of precursor ions provide better MS/MS fragmentation yields at lower collision energies (CE) during collision induced dissociation (CID).^24^ This is essential, as both the glycan moiety and the peptide moiety of glycopeptides need to yield sufficient fragment ions for an unambiguous characterization. When employing proteomics-like CID conditions on glycopeptides with low charge states, typically only glycan fragmentation is observed. Fragmentation of the peptide bond rarely yields sufficient b- and y-ions for unambiguous characterization of the peptide moiety. Potential solutions are the use of advanced fragmentation techniques like electron-based dissociation (ExD), that produce c- and z-type peptide fragment ions while retaining the intact glycan moiety.^25-26^ However, these techniques are not widely available. More commonly used CID instruments can overcome this obstacle by using higher collision energies to increase the yield of peptide fragment ions or performing CID stepping to fragment the glycan moiety at lower collision energies and the peptide moiety at high collision energies.^24, 27^

Another advancement for the holistic detection of glycopeptides in mixtures is the use of ion mobility mass spectrometry.^28^ Ion mobility is able to separate ions based on their size and shape in the gas phase and can provide and extra dimension of separation. One ion mobility technique is trapped ion mobility spectrometry (TIMS) in which ions are separated by finding an equilibrium between a constant gas flow and an electric field which allows them to be stored at different spatial positions inside the TIMS funnel. Ions can subsequently be eluted by lowering the electric field potential. In the timsTOF Pro and newer models (Bruker Daltonics), this technology enables parallel accumulation serial fragmentation (PASEF) for fast and sensitive fragmentation of orthogonally isolated precursor ions by mass-to-charge (*m/z*) and ion mobility. For the field of proteomics, this technology has already demonstrated its use by dramatically increasing the number of peptide identifications in bottom-up proteomics setups.^29^ Interestingly, the orthogonal precursor selection allows to focus the MS/MS spectra generation on glycopeptides rather than peptides as size and shape generally differs.^28^ Although these advancements in the detection of glycopeptides have been made, a combined methodology leveraging them in one workflow is missing. Hence, we established an integrated glycoproteomics workflow on the timsTOF Pro 2 that utilizes PASEF as well as the increased ionization efficiency of glycopeptides with DEN. We developed our glycoproteomics method starting from default settings for bottom-up proteomics and compared performance between glycopeptide and peptide identification to identify crucial parameters for glycoproteomic measurements.

We optimized electrospray conditions, ion optics, MS/MS precursor selection and collision energies to increase glycopeptide identification for enriched glycopeptides from tryptic human plasma digests. Our optimized method enabled identification of ∼1100-1500 glycopeptides using LC gradient times as short as 15 to 30 minutes, respectively, which enables comprehensive glycoproteome coverage at considerable sample throughput for (pre-)clinical applications.

## EXPERIMENTAL SECTION

### Tryptic digestion and glycopeptide enrichment from human plasma samples

Plasma samples of healthy donors were obtained from the Sanquin blood bank (Nijmegen, Netherlands) according to their protocols of informed consent. Samples from 5 individuals were pooled and preparation was performed as described previously.^4^ Ten microliters of human plasma were denatured with 10μL urea (8 M urea, 10 mM Tris-HCl, pH 8.0) and reduced with 15 μL of 10 mM dithiothreitol for 30 min at room temperature. Alkylation was performed by addition of 15 μL of 50 mM 2-chloroacetamide and incubation in the dark for 20 min at room temperature. Proteins were first digested with LysC (1 μg LysC/50 μg protein) for 3 hours at room temperature. Subsequently, samples were diluted with three volumes of 50 mM ammonium bicarbonate buffer. A tryptic digest was performed overnight at 37 °C by the addition of 1 ug trypsin per 50 ug protein. Glycopeptides were enriched using Sepharose CL-4B beads (Merck). 100 μL of slurry were added to a well of a 0.20 μm pore size 96 multi well filter plate (AcroPrep Advance, VWR). The beads were washed three times with 20% ethanol and 83% acetonitrile. After applying the digested sample, the plate was incubated for 20 min at room temperature while shaking. The filter plate was centrifuged to remove the supernatant and beads were washed three times with 83% ACN and three times with 83% ACN with 0.1% trifluoroacetic acid (TFA). Glycopeptides were eluted with 50 μL water.

### LC-MS/MS analysis

The samples were analyzed employing a nanoElute liquid chromatography system (Bruker Daltonics) connected to a timsTOF Pro 2 instrument (Bruker Daltonics). Unless otherwise stated, a CaptiveSprayer nanoflow electrospray ionization source with dopant-enriched nitrogen gas was used via the nanobooster. Acetonitrile was used as solvent. Separation of peptides and glycopeptides was achieved on an Elute Fifteen C18 reversed-phase column (0.075 mm ID × 150 mm length, 1.9 µm particles, 120 Å pore size) operated at 45 °C. The elution gradient for most optimization steps consisted of a linear increase from 7% to 45% acetonitrile in 0.1% formic acid (FA) and 0.02% TFA over 15 minutes, with a flow rate of 500 nL/min. More information on the used gradients can be found in supplementary table S1. Mass spectrometry measurements were conducted in positive ionization mode with a capillary voltage of 1500 V and, if used, a nanobooster gas pressure of 0.2 bar N_2_. MS conditions were optimized starting from default setting for proteomics measurements provided by Bruker Daltonics and adjusted to facilitate glycopeptide identification. The MS conditions of the proteomics method and optimized glycoproteomics methods can be found in supplementary table S2. All measurements were performed in duplicates.

### Optimization strategy

We optimized several acquisition parameters to establish an integrated glycoproteomics method on the timsTOF Pro 2. First, we optimized collision energies to achieve ideal MS/MS fragmentation. Subsequently, we optimized ion optics setting to enhance transmission of glycopeptides. PASEF conditions, namely the target intensity, TIMS isolation width (also referred to as measuring time) and the polygon region for PASEF precursor selection were adjusted. Finally, we evaluated the effect of different LC gradients and benchmarked our optimized method with and without the use of DEN. And overview of all used MS acquisition parameters for each optimization step can be found in supplementary table S3. All MS data is deposited in ProteomeXchange repository under the identifier PXD047898.^30^

### Data interpretation

Glycopeptides were identified using MSFragger Glyco. MSFragger searches were performed using fragpipe 15.0, MSFragger 3.4, and philosopher 4.1.1. The glyco-N-HCD search parameters were: 20 ppm mass tolerance with an isotope error of 0–2, semi-tryptic enzyme specificity, peptide length 5–50 amino acids, and 600–20,000 m/z range. Carbamidomethylation at cysteine residues was set as fixed modification whereas methionine oxidation and N-terminal ammonia loss were set as variable modifications. Human secreted protein reference sequence database was downloaded from Uniprot (downloaded on 2021.11.22) and glycan mass offsets were extracted for unique compositions in the GlyGen glycan reference database (downloaded on 22.4.2022).^31-32^ The FDR was set to 1% at PSM, peptide, protein, and glycan levels. DataAnalysis v5.3 (Bruker Daltonics) was used for raw data analysis. The used software MSFragger Glyco identifies glycan compositions based on mass offsets. With this approach, high fragmentation of the peptide moiety yields optimal results. Other software tools that take glycan fragmentation into account can benefit from collision energy stepping.^24-25, 33^ We therefore provide the optimized instrument method both with and without TIMS stepping in the ProteomeXchange repository under the identifier PXD047898.

Data was visualized using Python 3.10.10 with the packages Matplotlib 3.7.1, Scipy 1.10.1, Numpy 1.24.3 and Pandas 1.5.3. All data points represent the average of two replicate measurements. Peptide and glycopeptide images were created using PyMOL 2.5.5.

## RESULTS AND DISCUSSION

To develop a generic analytical PASEF workflow for holistic glycoproteomics, we used enriched glycopeptides from human plasma samples. Although our methodology can be applied to various sample types, enriched glycopeptides from human plasma not only represent a complex mixture with heterogenous glycopeptides but are also biologically and clinically relevant. The enriched glycopeptides were subjected to LC-MS/MS measurements, starting with the default parameters for proteomics on the timsTOF platforms (for proteomics parameters see supplementary table S2). We subsequently optimized collision energies for glycopeptide fragmentation, electrospray source conditions and ion optics, PASEF settings and gradient length. To verify the outcome of our optimization steps, we evaluated results based on the number of identified glycopeptides in relation to the number of peptide identifications. MSFragger Glyco software was used for (glyco)peptide identification, which identifies the peptide moiety of glycopeptides using a modified open mass search strategy whereas glycan moiety compositions are identified based on mass offset and limited glycan fragmentation data in combination with reference glycan compositions.^34^ Finally, we benchmarked our optimized method against the standard proteomics method using acetonitrile-enriched nitrogen gas to enhance the ionization efficiency of glycopeptides. A schematic overview of the instrument configuration and conceptual optimization steps is shown in Figure 1.

**Figure 1.**
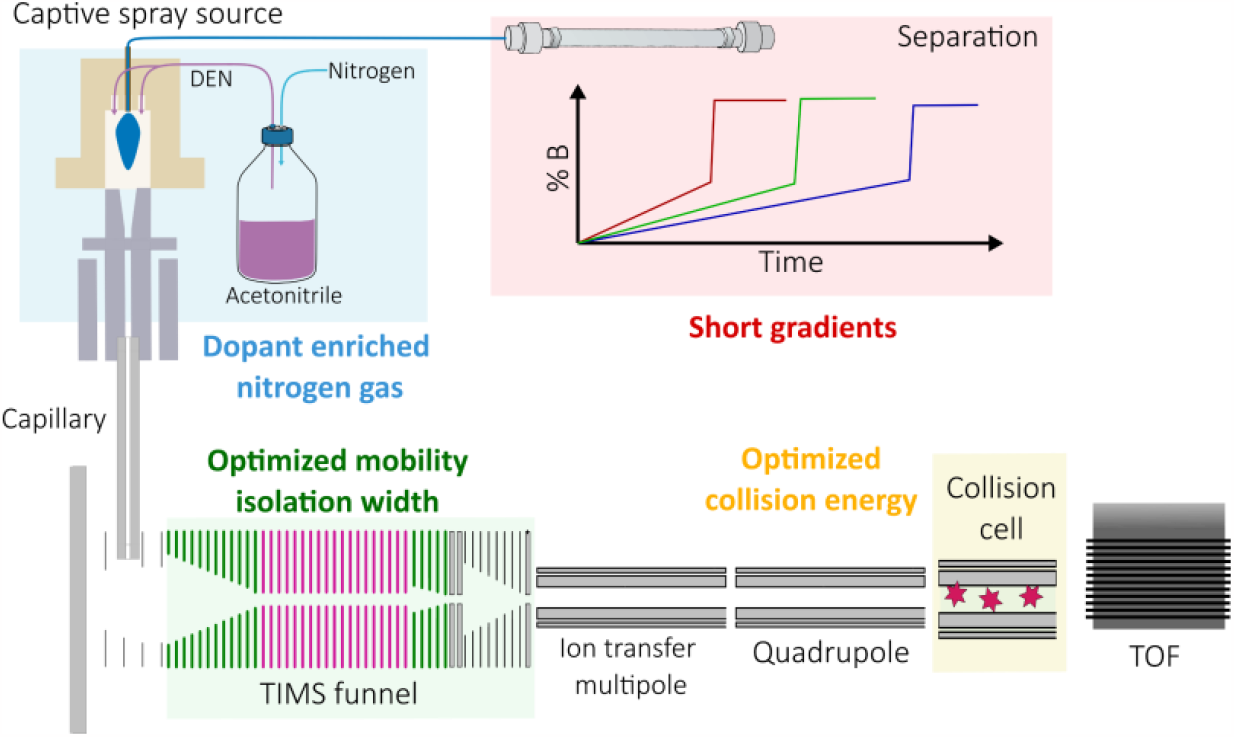
Optimization scheme. Our methodology integrates the advantages of DEN, optimized PASEF settings with ideal mobility isolation width and collision energies to increase glycopeptide identification with short LC run times of 15 to 30 minutes.

### Optimizing CID energies

The identification of glycopeptides by tandem mass spectrometry poses a significant challenge due to the need for optimal fragmentation of both the peptide moiety and the intrinsically different glycan moiety. Conventional shotgun proteomics CID energies typically yield predominant glycan fragmentation with insufficient peptide fragment ions to confidently identify the glycopeptide.^35^ To address this limitation, we systematically adjusted collision energies to enhance the yield of peptide backbone fragments in our approach.

The timsTOF Pro 2 instruments enable collision energy scaling based on precursor ion mobility values. At lower ion mobility values, higher charge states and smaller glycopeptides are expected, requiring less energy to yield peptide backbone fragments. Conversely, higher mobility values indicated lower charge states and larger glycopeptides. Starting from the default proteomics collision energy slope we incrementally increased the slope by steps of +10% up to +60% as seen in Figure 2, A. An almost linear increase in the number of unique glycopeptide identifications with increasing collision energies was observed (Figure 2, B). In contrast, the number of peptide identifications decreased at higher collision energies. Furthermore, the optimal collision energy values were found to be dependent on the precursor charge state. For example, glycopeptides with a charge state of +5 did not benefit from increasing the collision energies above +30% as illustrated in Figure 2, C. As our final optimized glycoproteomics method employs acetonitrile-enriched nitrogen gas, resulting in an overall charge state increase of glycopeptides, we did not increase collision energies any further and continued our optimization steps with a collision energy increase of +50% compared to default proteomics settings. Additionally, it’s worth noting that collision energies in most timsTOF Pro instruments are limited to 100 eV.

**Figure 2.**
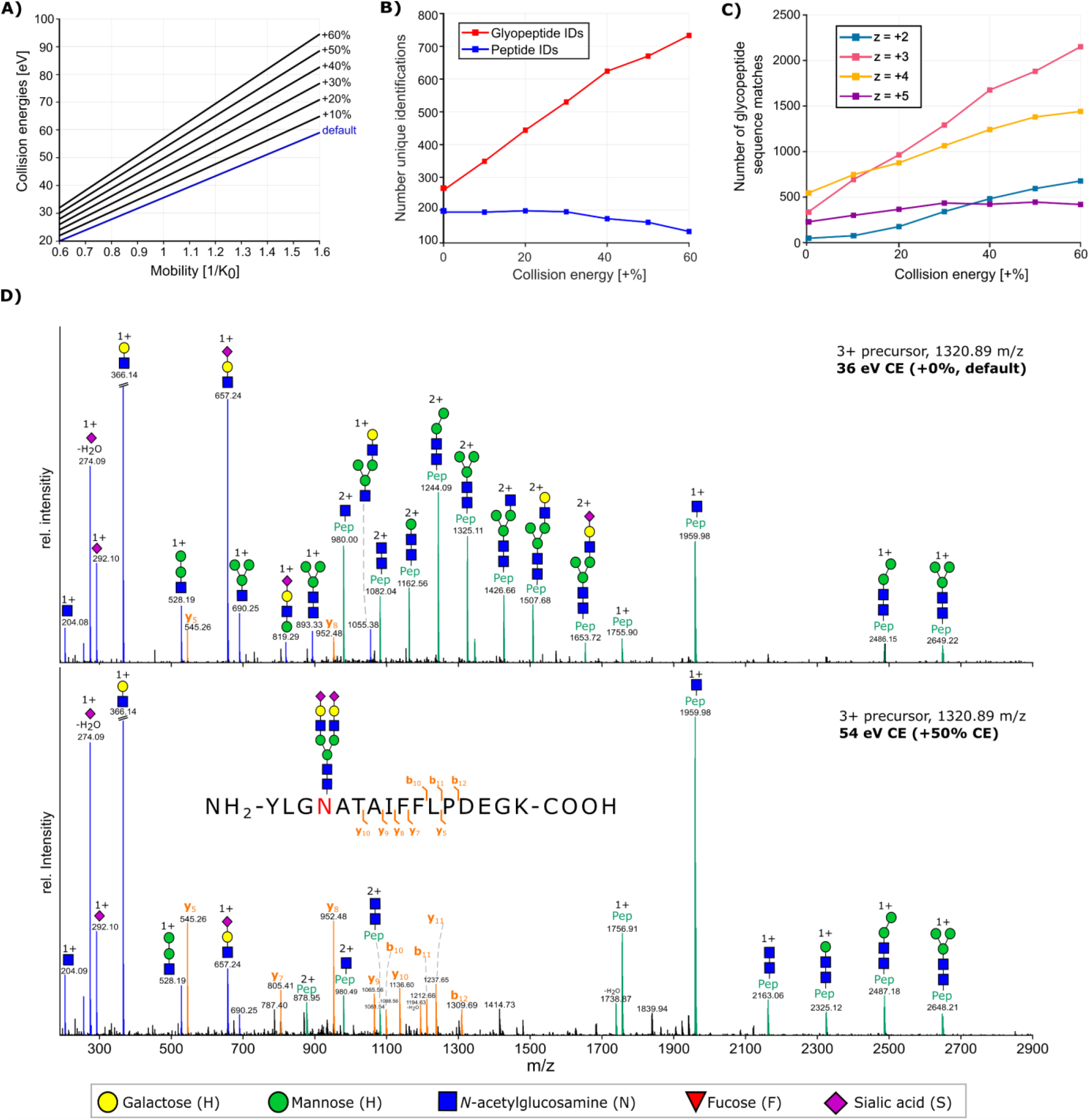
Collision energy optimization. (A) collision energies are set as a function of ion mobility and increase linear with higher mobility values. The slope and offset were increased in steps of 10% compared to the proteomics default settings to optimize collision energies for glycopeptide fragmentation. (B) Increasing the collision energies led to an increase in unique glycopeptide identifications from less than 300 to over 700 at higher CE values while peptide identifications decreased. (C) Ideal CE values depended on precursor charge state. Precursors with a lower charge state (+2, +3) benefited more from increased CE values compared to highly charge precursors (+4, +5). (D) The exemplary MS/MS spectrum of a +3 glycopeptide with standard proteomics CE values shows mainly the fragmentation of the glycan moiety as oxonium ions (blue) or glycan fragments with intact peptide residue (green). At higher CE values (+50%), more b- and y-ion fragments of the peptide backbone (orange) are observed to support peptide moiety identification by protein sequence database search algorithms. The annotation of glycan fragments represents only one possibility. Other isomers are plausible.

### Improving ion transmission

Optimizing the collision energies dramatically increased the number of unique glycopeptide identifications from under 300 identifications to over 700 identifications. To further increase identifications, we continued with optimizing ion optics, namely the prepulse storage and transfer time which are responsible for the transmission of ions from the collision cell to the TOF orthogonal accelerator.

Furthermore, we optimized the collision cell RF. Ramping of the prepulse storage and transfer time did not result in a strong increase of glycopeptide identifications over peptide identifications (supplementary Figure S1). In contrast, we found that the collision cell RF has a strong influence on glycopeptide identification and that increased Vpp values promote the identification of glycopeptides while decreasing the identification of peptides (supplementary Figure S1). With increased RF Vpp, the kinetic energy of ions is increased, enabling them to maintain their trajectory through the collision cell, thus improving the transmission of ions with higher *m/z* values.^36-37^ This is crucial as glycopeptides show y-ion fragments at high *m/z* values due to fragment ions containing the whole peptide backbone and different glycan fragments at reduced charge states. For glycopeptide identification, determining the correct peptide mass based on the Y0 “peptide-only” or Y1 “peptide+HexNAc” fragment is essential (see annotation of MS/MS spectra in Figure 2, D).^24, 34^ These Y-ions are observed at higher *m/z* values compared to peptide fragment ions. Hence, we increased the collision cell RF from 1500 eV to 1700 eV.

### Optimizing PASEF settings

One of the key features of the timsTOF platform, which provides high sensitivity and fast MS/MS acquisition with simultaneous ion mobility separation, is parallel accumulation and serial fragmentation (PASEF) of ions.^38-39^After drastically increasing glycopeptide identification by adjusting collision energies, we evaluated PASEF parameters for optimal MS/MS data generation. First, we tested the effect of the target intensity for MS/MS acquisition as we expected that higher target intensities would lead to an increased intensity and signal-to-noise ratio of peptide backbone ions. However, increasing the target intensity in a range from 20.000 counts to 150.000 counts did not result in an increase of glycopeptide identifications (see supplementary Figure S2). Next, we focused on the tims isolation width as PASEF precursor ion selection is based on *m/z* values and mobility. As ions are sequentially released from the tims funnel, the quadrupole is isolating precursor ions by quickly switching between different *m/z* positions while achieving MS/MS acquisition rates of up to 110 Hz in the timsTOF Pro 2. Considering typical ion mobility peak widths of peptides, the quadrupole isolation time is set between 2.5 and 4.0 ms which can theoretically result in the selection of 240 to 400 precursor ions per second.^39^ For our measurements, we chose a ramp time of 100 ms with 100% duty cycle. In the default proteomics method, the mobility range is set from 0.6 to 1.6 1/K_0_ [V·s/cm^2^] and the tims isolation width is set to 2.75 ms. This results in a mobility isolation window of ±0.028 1/K_0_ [V·s/cm^2^]. Isolation widths of 2.75 ms are ideal for peptides, where typical half-widths of 1 ms are observed under similar conditions.^39^ However, this isolation width might not be ideal for glycopeptides as the glycan moiety not only consist of complex isomers but also multiple conformations for each isomer.^40^ Hence, we tested tims isolation widths of 2.75, 5.00, 7.50 and 10.0 ms to determine the effect on glycopeptide identification rates. Figure 3A shows the structure of one selected peptide and one selected glycopeptide to demonstrate typical differences in size but also rotatable bonds in peptides and glycopeptides. Mobilograms for both the peptide and glycopeptide are shown in Figure 3B. The TIMS isolation width of 2.75 ms is sufficient to capture the entire mobility peak for the peptide, however, a broader isolation width is needed for glycopeptides to avoid ion loss during MS/MS isolation. When ramping the isolation width from 2.75 to 10 ms, the number of unique glycopeptide identifications increases with isolation widths up to 7.5 ms before decreasing at 10 ms (Figure 3C). For peptides, an increase in isolation width decreases the number of identifications. This could be due to a reduction in MS/MS detection of unique precursor ions, as fewer precursor ions are selected per TIMS ramp.

**Figure 3.**
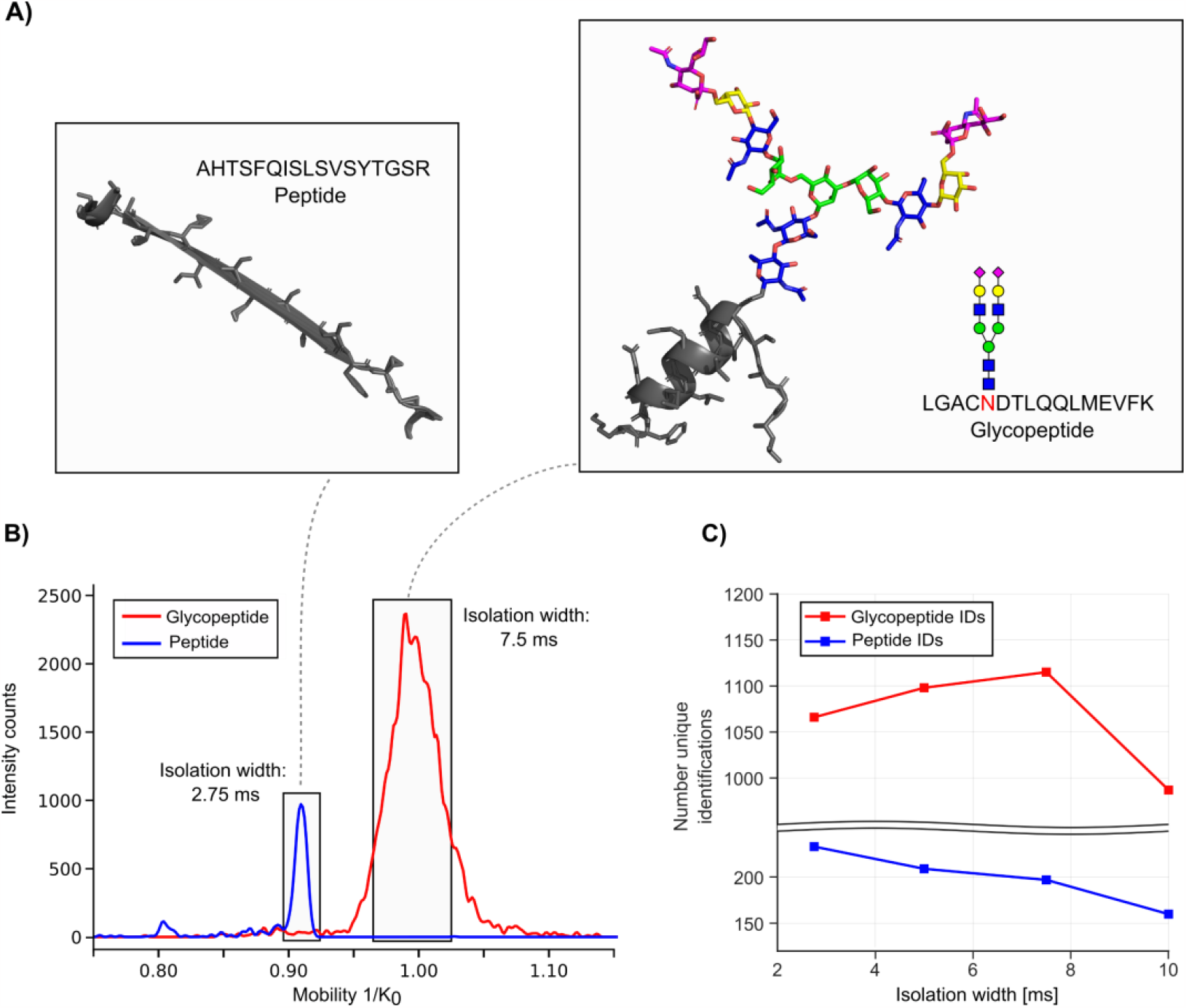
TIMS isolation width optimization. (A) Structure of one selected peptide (pdb entry 7VON, alpha-2-macroglobulin) and one selected glycopeptide (pdb entry 1ATH with attached glycan, human antithrombin III). (B) Mobilogram of the selected peptide (blue, sequence AHTSFQISLSVSYTGSR) and glycopeptide (red, sequence LGAC**N**DTLQQLMEVFK decorated with the glycan composition H5N4S2). The standard TIMS isolation width of 2.75 ms is ideal for the isolation of the peptide, however, a much larger isolation with of 7.5 ms is needed for the glycopeptide due to a broader mobility distribution. (C) Number of peptide and glycopeptide identifications with increasing TIMS isolation width.

In addition to increasing the TIMS isolation width, we specifically selected a mass and mobility region that favored the selection of glycopeptides over peptides as MS/MS precursor. As demonstrated before, glycopeptides show a distinct difference in mass and ion mobility and selecting this region of interest by defining a polygon for MS/MS precursor selection can increase the number of MS/MS spectra of glycopeptides.^28^ The polygon used in the proteomics default setting and our revised polygon can be found in supplementary Figure S3.

### Influence of LC-gradient length

High sample throughput is not only a crucial parameter in bottom-up proteomics but also essential for glycoproteomics to enable experiments with large sample cohorts or routine testing in clinical setups. To achieve higher throughput, liquid chromatography gradients can be reduced if the peak capacity is still sufficient and MS/MS sampling is fast.^41^ The timsTOF Pro 2 offers additional separation power in the mobility dimension as well as fast MS/MS acquisition rates to sample LC peaks with small peak widths, therefore facilitating high identification rates with short LC gradients. To determine how gradient length affects glycopeptide identification on our 15 cm reversed-phase nanoflow column, we tested different gradient lengths from 5 to 60 minutes (Figure 4). Information on the gradient conditions can be found in supplementary Table S1.

**Figure 4.**
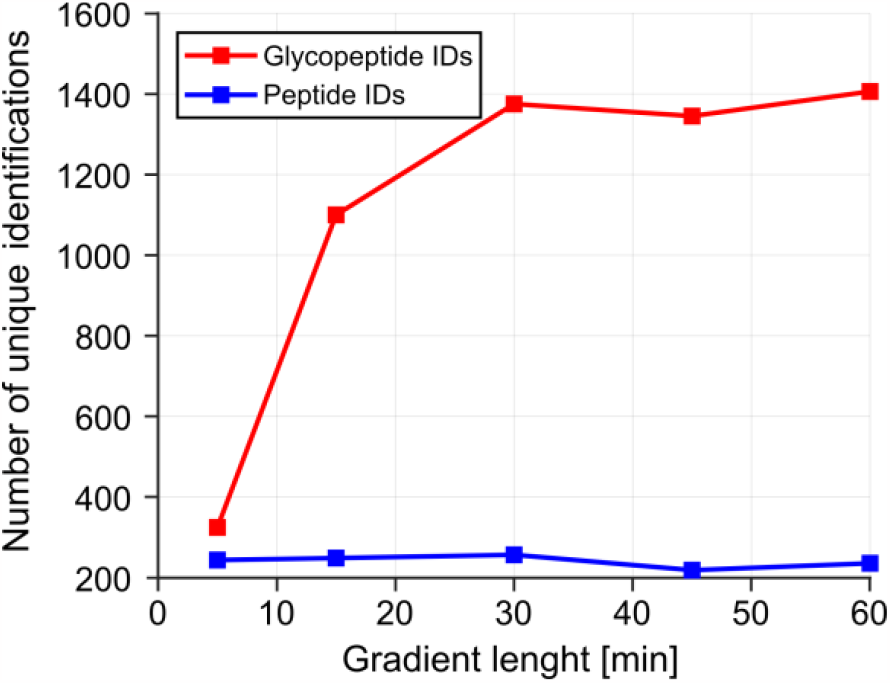
Influence of gradient length on the number of unique peptides (blue) and glycopeptides (red).

Unique glycopeptide identifications drastically increased when increasing between gradient times of 5 to 15 minutes from over 300 to almost 1200 identifications. Longer gradient times of 30 minutes still showed an improvement in glycopeptide identifications, however, the number of identifications stagnated with gradient times over 30 minutes. Hence, for the following measurements, a gradient length of 30 minutes was selected.

### Benchmarking of optimized conditions under use of dopant-enriched nitrogen gas

After combining optimized parameters, we benchmarked our glycoproteomics method against the bottom-up proteomics default method with and without the use of DEN. As reported previously, the use of DEN can drastically increase ionization efficiencies of polar compounds such as glycans, glycoproteins, and glycopeptides.^20-23^ Hence, we evaluated how much our use of acetonitrile as DEN influences glycopeptide identification rates under proteomics settings as well as with optimized glycoproteomics settings. As expected, the use of DEN strongly influences the number of glycopeptide identifications for both the default proteomics method and the developed glycoproteomics method as seen in

We observed that not only glycopeptide identifications increase with the use of DEN, but also the number of peptide identifications. Still, the use of DEN seems to have a stronger effect on glycopeptides compared to peptides, as shown by the fourfold increase in glycopeptide identifications under proteomics conditions and the increase in identifications from 960 to almost 1500 under glycoproteomics conditions (Figure 5).

**Figure 5.**
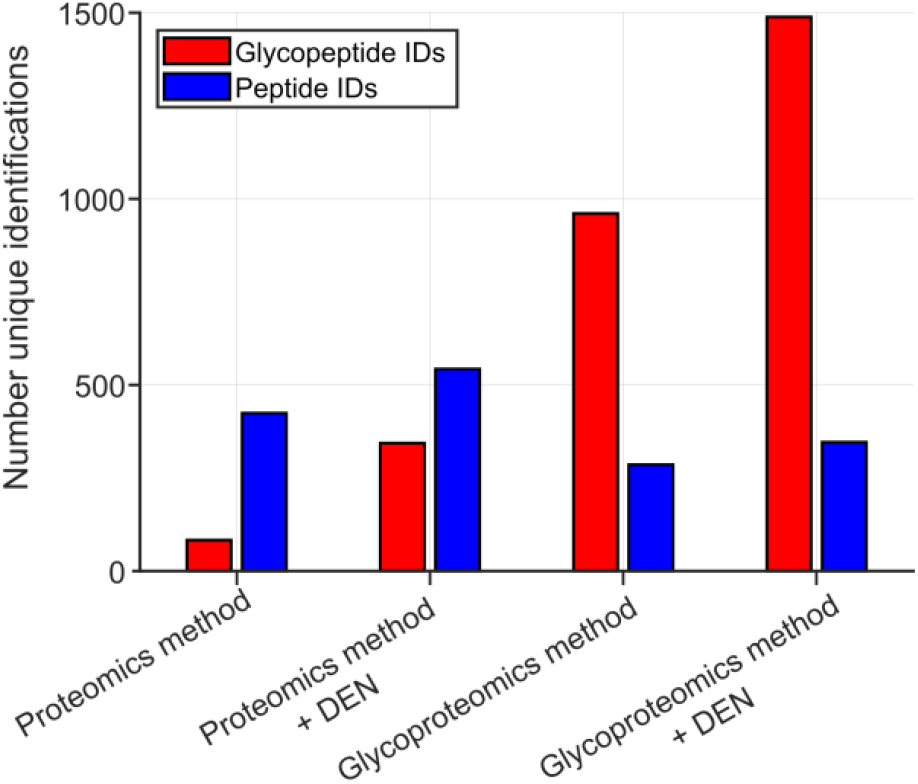
Benchmarking of glycoproteomics method against default proteomics settings employing a 30 minutes gradient with and without the use of DEN.

It is worth mentioning that when DEN was used, the mobility values for peptides and glycopeptides experienced a small shift towards higher mobility which can be seen in supplementary Figure S3. To evaluate the significances of shifting mobility values with the use of DEN, we measured the shift in mobility for one calibrant ion with different nanobooster fill levels as seen in supplementary Figure S4. A slight shift in mobility was observed, however, a regular refilling of the nanobooster with acetonitrile enabled us to obtain stable results during glycoproteomics measurements.

In addition, we were interested to see how charge states are influenced by using DEN. Hence, we plotted the glycopeptide spectrum matches per charge state in the *m/z* and mobility dimension for all four conditions (Figure 6). Without the use of DEN, we dominantly observe glycopeptide spectrum matches with a charge state of +3. This distribution shifts towards an almost equal occurrence of charge states of +3 and +4 with DEN both under default proteomics conditions and when using the optimized glycoproteomics method (Figure 6 and supplementary Figure S5). Remarkably, the number of glycopeptide spectrum matches with a charge state of +2 is low when using the default proteomics method with and without DEN (Figure 6). We conclude that this is caused mainly by the low collision energies of the proteomics method, which do not allow sufficient fragmentation of glycopeptide ions with low charge states (see also Figure 2, panel C).

**Figure 6.**
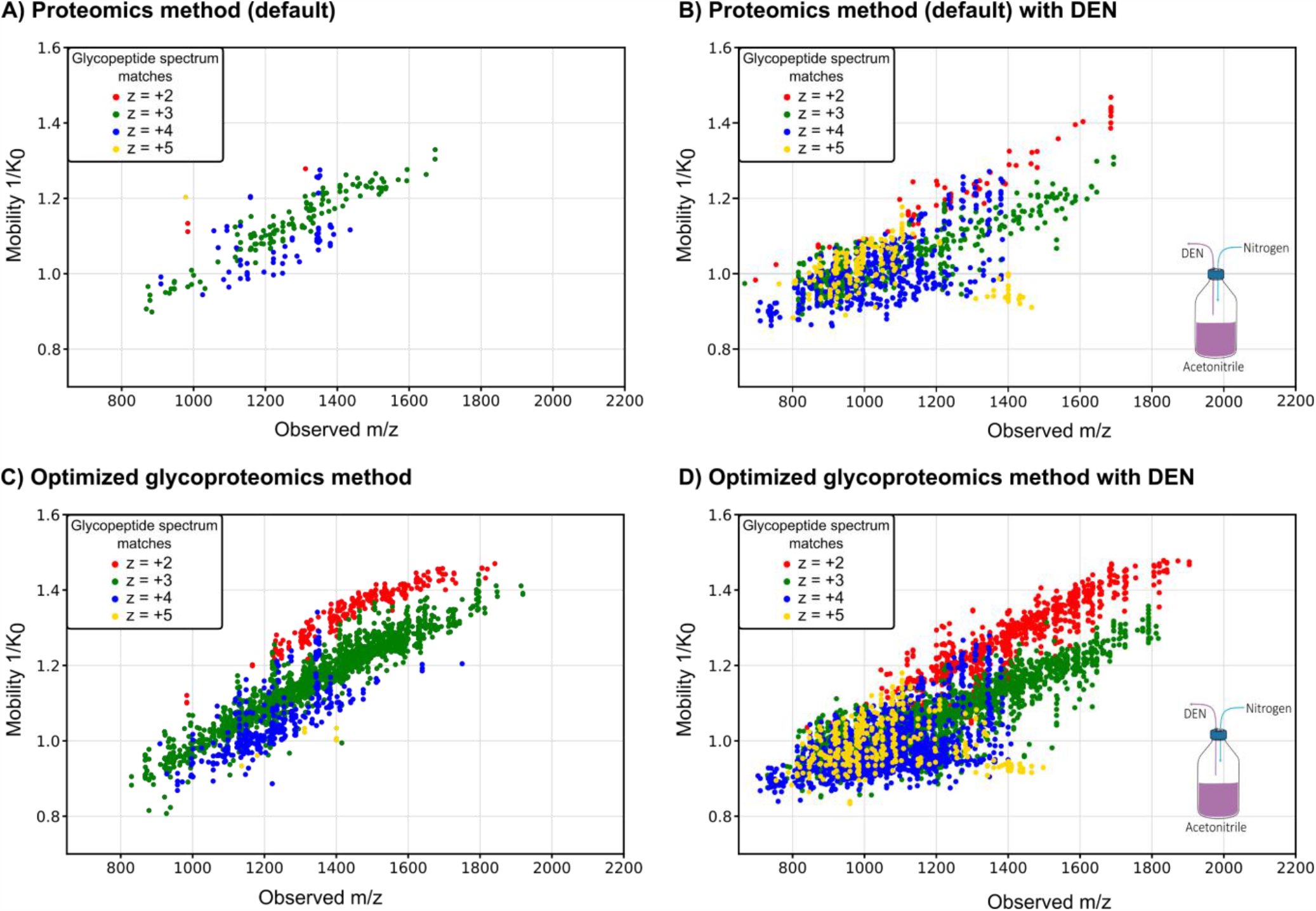
Charge state distribution of glycopeptide sequence matches with and without the use of DEN. (A) Using standard proteomics condition without DEN results in a low number of glycopeptide spectrum matches. (C) The optimized glycoproteomics method drastically increases the number of glycopeptide spectrum matches. For both conditions, the dominant charge state is +3. (B and D) When using DEN, the number of glycopeptide spectrum matches drastically increases for both methods. In addition, more ions with a charge state of +4 and +5 are observed.

## CONCLUSION

In this work, we have developed an optimal integrated PASEF workflow for glycoproteomics measurements. By optimizing multiple parameters, including collision energies (CE), ion optics, and ion mobility settings, we established a robust workflow for the holistic identification of glycopeptides. Our results show that the use of dopant-enriched nitrogen (DEN) gas and optimization of collision energies have a significant impact on glycopeptide identification rates. Unlike most approaches that optimize collision energies or report the benefits of DEN on a limited number of glycopeptides, our study provides a broader perspective by evaluating a larger number of glycopeptides.^20, 28^ However, it must be considered that the ideal collision energy values reported here are advantageous for the used glycan offset search strategy and that other data interpretation platforms may require slightly different settings to target the glycan moiety of glycopeptides.

By comparing peptide and glycopeptide identification rates, we demonstrate the need to adjust parameters based on the molecular characteristics of the analytes. This emphasizes the inherent complexity of glycoproteomics and underlines that a one-size-fits-all approach is not applicable to obtain optimal results. Our adjusted methodology, which integrates optimized collision energies, PASEF settings, and DEN, provides a high-throughput approach suitable for glycopeptide analysis in complex biological samples on the widely used timsTOF platform. Thus, this work contributes to the growing field of glycoproteomics and paves the way for improved characterization of post-translational modifications in complex biological samples, enabling broad applications in both research and (pre-) clinical settings.

## ASSOCIATED CONTENT

### Supporting Information

The supporting information contains Figures S1 to S5 and Tables S1 to S3 for additional information on optimization steps.

## AUTHOR INFORMATION

### Author Contributions

The manuscript was written through contributions of all authors. All authors have given approval to the final version of the manuscript.

## ACKNOWLEDGMENT

The collaboration project EnFORCE (Enabling Functional Omics in Routine Clinical Environments, LSHM21032) is co-funded by the PPP Allowance made available by Health∼Holland, Top Sector Life Sciences & Health, to stimulate public–private partnerships. This research was financially supported by infrastructure support from the ZonMw Medium Investment Grant 40-00506-98-9001, the Radboud Consortium for Glycoscience, an interfaculty grant from Radboud University, and EUROGLYCAN-omics (ERARE18-117) by ZonMw (90030376501), under the frame of E-Rare-3, the ERA-Net for Research on Rare Diseases and is part of the Netherlands X-omics Initiative, partially funded by NWO (project 184.034.019).

## Notes

### Competing Interest Statement

The authors have declared no competing interest.

### Summary of Updates

Glycoproteins play important roles in numerous physiological processes and are often implicated in disease. Analysis of site-specific protein glycobiology through glycoproteomics is evolving rapidly in recent years thanks to hardware and software innovations. Particularly, the introduction of Parallel Accumulation Serial Fragmentation (PASEF) on hybrid trapped ion mobility time-of-flight mass spectrometry instruments combined deep proteome sequencing with separation of (near-)isobaric precursor ions or converging isotope envelopes through ion mobility separation. However, reported use of PASEF in integrated glycoproteomics workflows to comprehensively capture the glycoproteome is still limited. To this end, we devel-oped an integrated methodology using the timsTOF Pro 2 to enhance N-glycopeptide identifications in complex mixtures. We systematically optimized the ion optics tuning, collision energies, mobility isolation width and the use of dopant-enriched nitrogen gas (DEN). Thus, we obtained a marked increase in unique glycopeptide identification rates compared to standard proteomics settings showcasing our results on a large set of glycopeptides. With short liquid chromatography gradients of 30 minutes, we increased the number of unique N-glycopeptide identifications in human plasma samples from around 100 identifications under standard proteomics condition to up to 1500 with our optimized glycoproteomics approach, highlight-ing the need for tailored optimizations to obtain comprehensive data.

