## Supplementary information for "Maximizing glycoproteomics results through an integrated PASEF workflow"

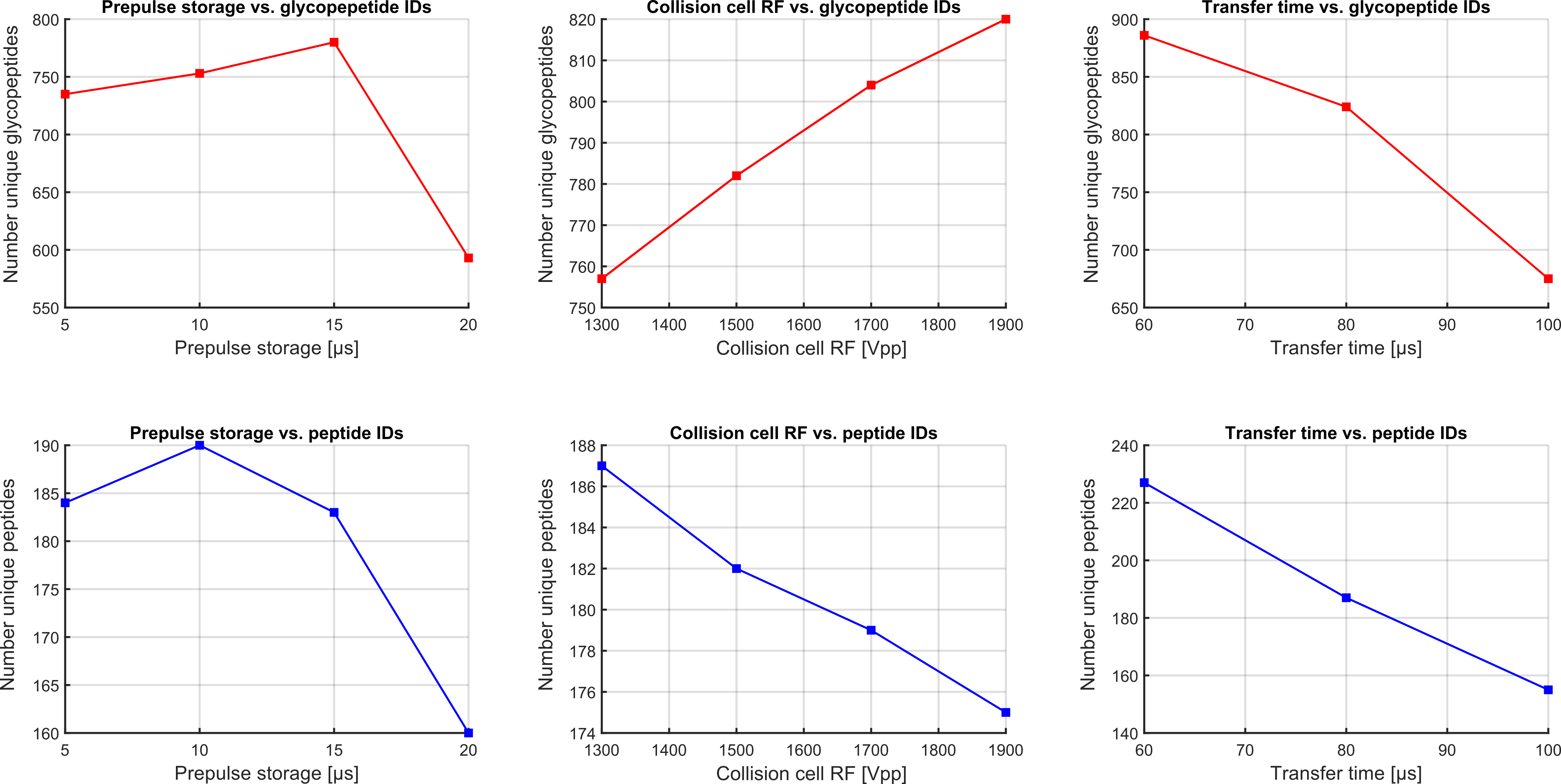
**Figure S1.** Tuning of ion optics for ideal transmission and identification of glycopeptides via optimization of prepulse storage time, collision cell RF and transfer time. Shown in top panels are glycopeptide IDs (red) whereas the bottom panels show peptide IDs (blue). Of particular interest is the collision cell RF. A higher collision cell RF is favorable for glycopeptide identifications. This is potentially due to the occurrence of glycopeptide fragment ions at higher m/z values caused by fragmentation of the glycan moiety with intact peptide moiety. These fragment ions, in particular the peptide+HexNAc peak, are relevant for the correct assignment of the peptide mass of glycopeptides and benefit from higher collision cell RF values. Although 1900 Vpp shows the highest number of glycopeptide IDs, we selected a value of 1700 Vpp for the following measurements since settings above 1700 Vpp led to a significant signal loss of low m/z oxonium ion intensities.

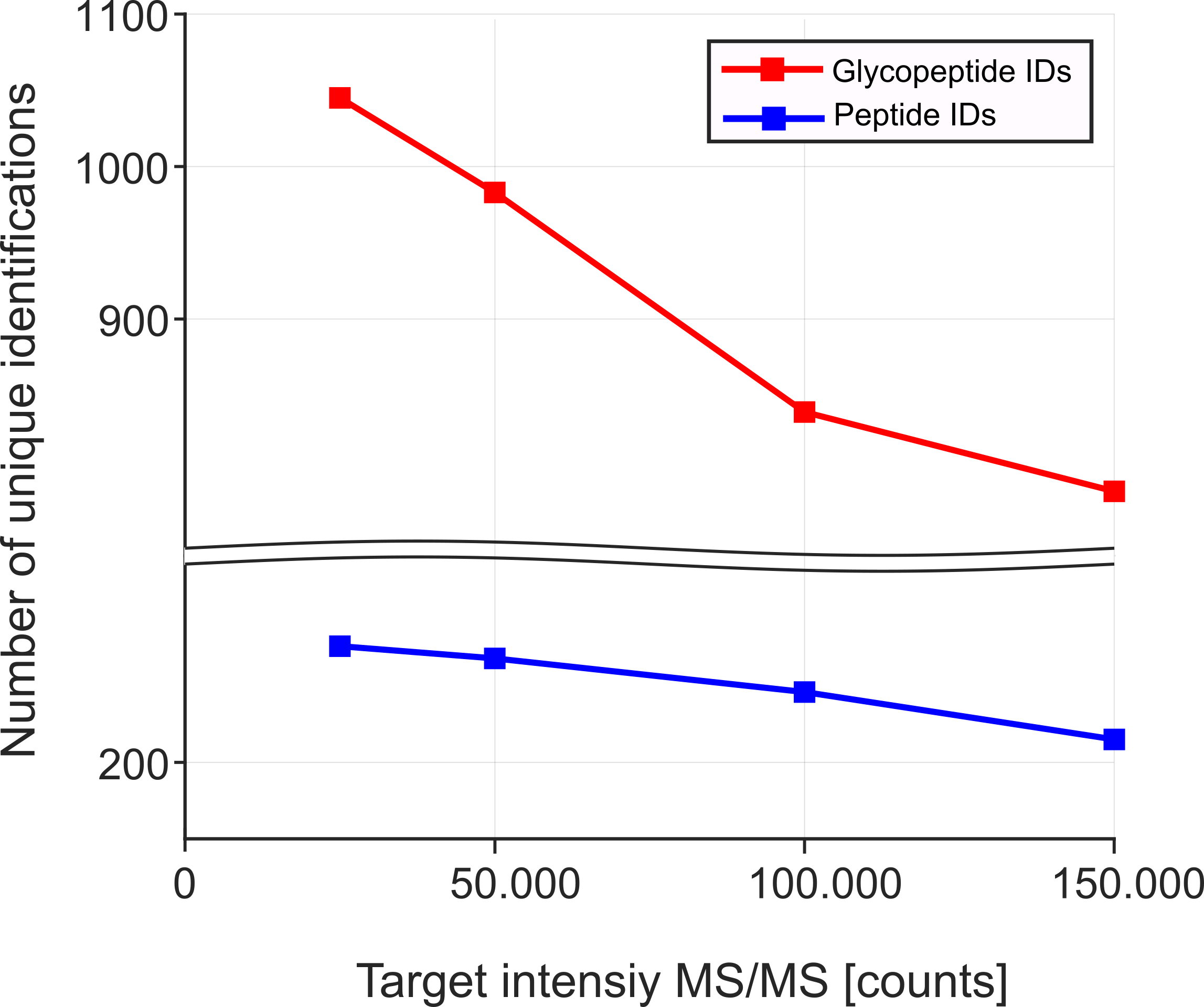

**Figure S2.** Optimization of target intensity. Both for peptides and glycopeptides, increasing the target intensity does not improve identification. For the developed glycoproteomics method, a target intensity of 25.000 counts was selected which is comparable to the target intensity of 20.000 in the default proteomics method.

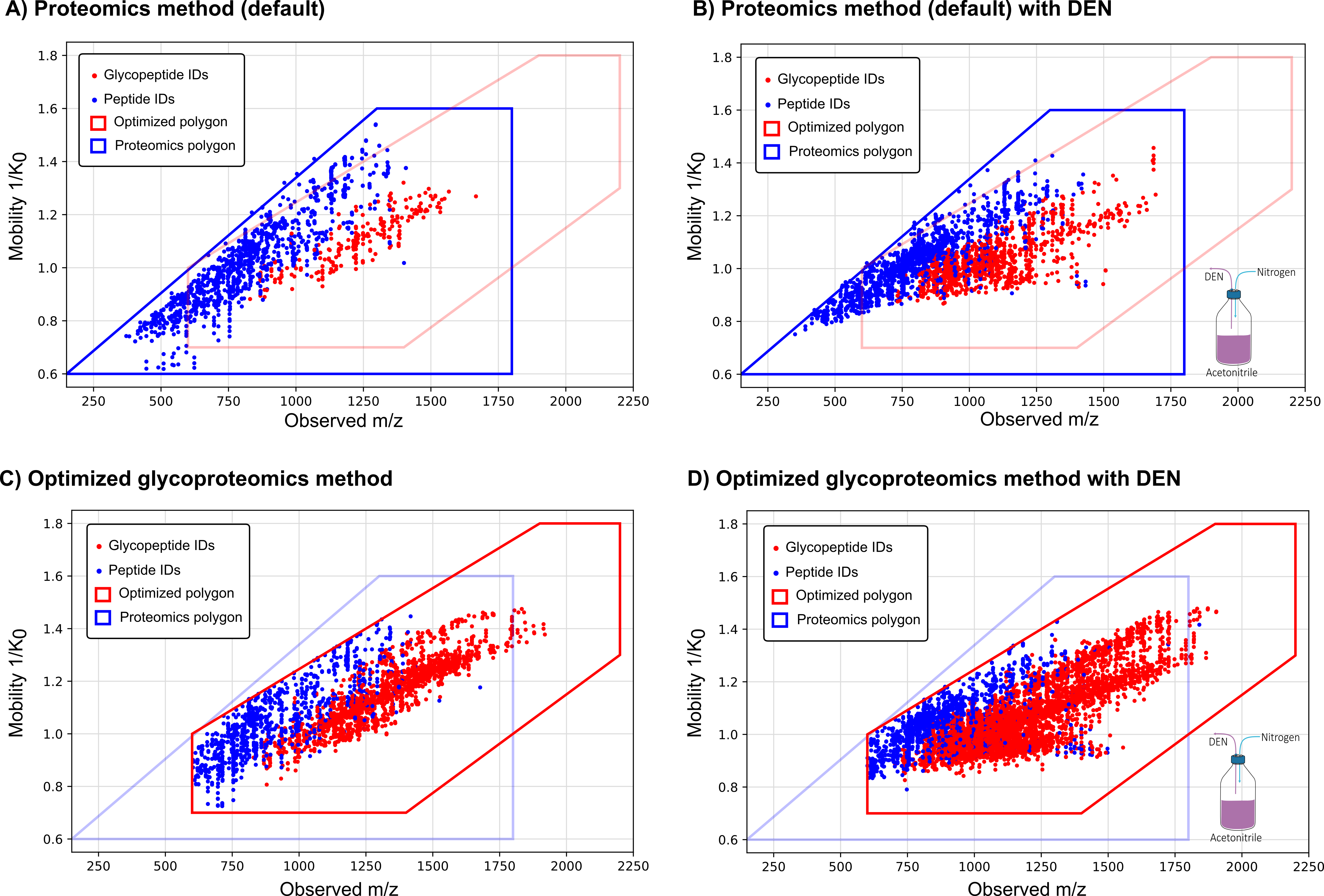

**Figure S3.** Glycopeptide and peptide IDs plotted by m/z vs. mobility value for proteomics method and optimized glycoproteomics method with and without the use of dopant enriched nitrogen gas via nanobooster. The default polygon used in the proteomics method is depicted in blue and the optimized polygon for glycopeptide identification is plotted in red. The proteomics polygon was used in A and B, the glycoproteomics polygon was used in C and D.

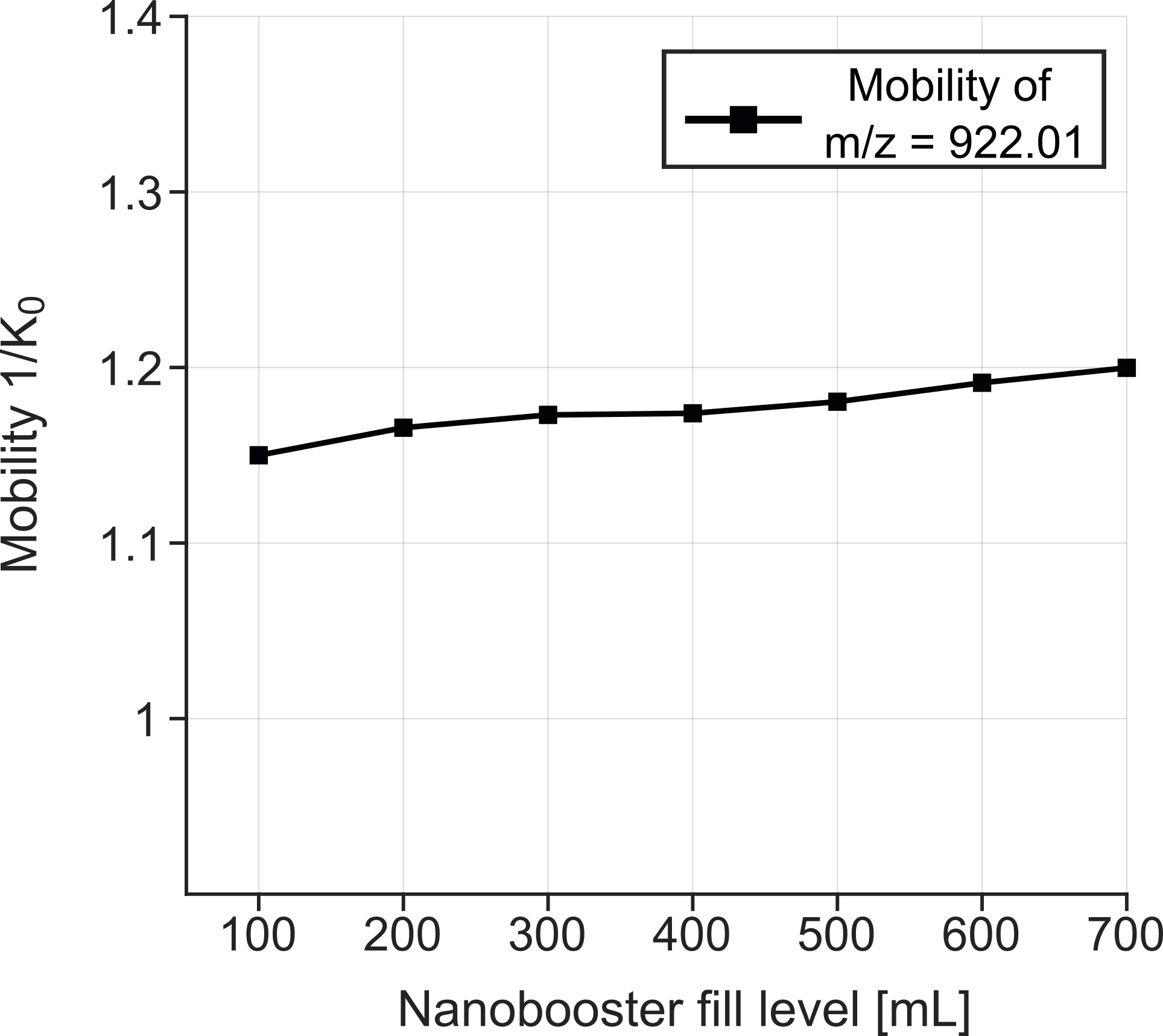

**Figure S4.** Change in mobility of one agilent tunemix (Agilent Technologies) signal at m/z = 922.01 with increasing fill levels of the nanobooster with acetonitrile. When using DEN, the mobility of the ions shifts as the collision gas density, flow rate and possibly temperature change. In addition, there is a slight shift in mobility over time as the solvent content in the nanoBooster decreases as shown here. This mobility shift can be well controlled if the nanoBooster is refilled regularly.

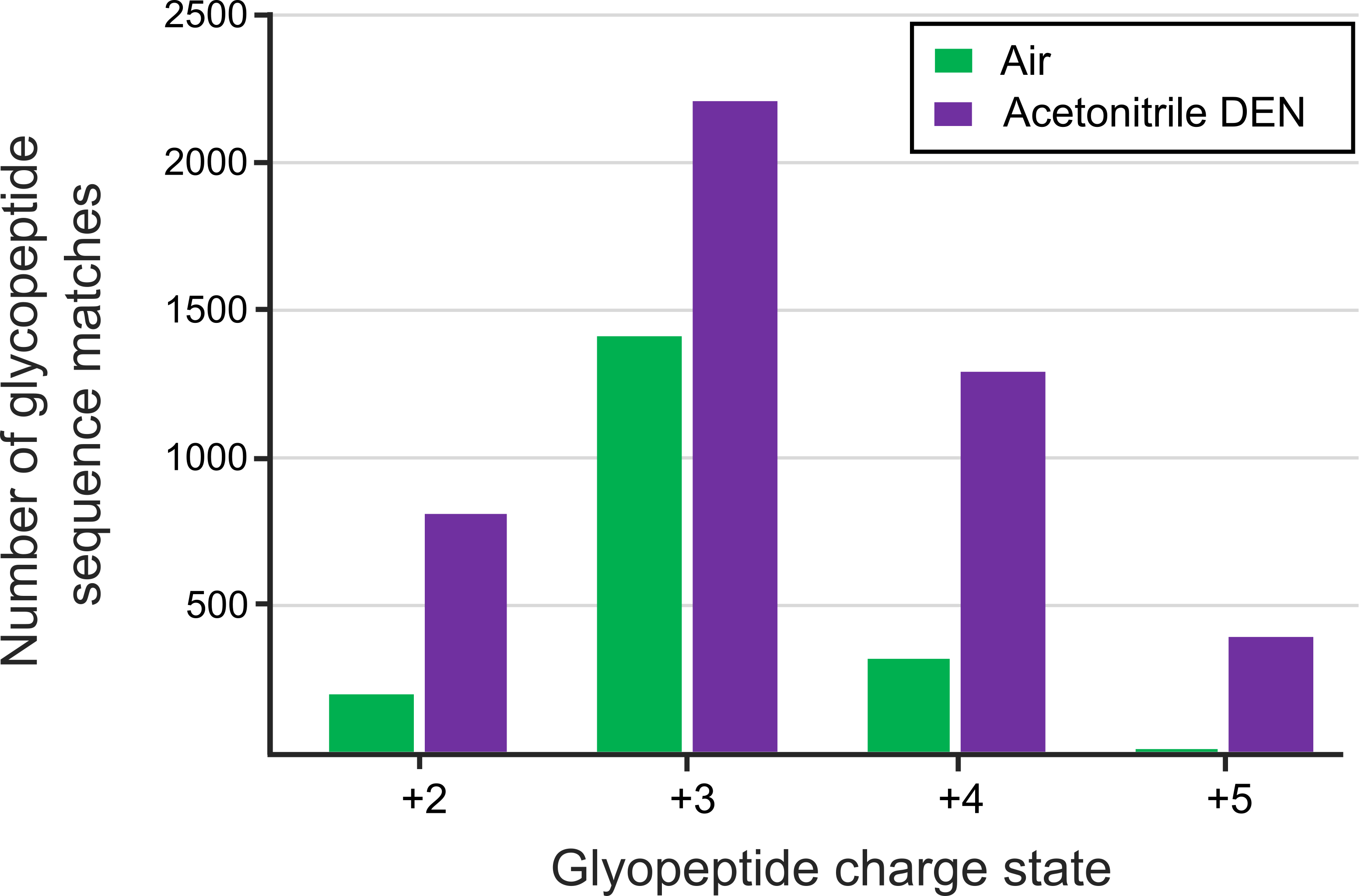

**Figure S5.** Charge state distribution of glycopeptides with and without acetonitrile-enriched nitrogen gas using the optimized glycoproteomics method. The number of glycopeptide sequence matches represents the average between two replicate measurements. A dominant charge state of +3 is observed without the use of DEN. When using DEN, the chare state distribution shift towards an almost equal amount of +3 and +4 precursor ions.

**Table S1.** Tested nanoLC gradients. Solvent A was composed of water with 0.1% FA and 0.02% TFA. Solvent B was composed of acetonitrile with 0.1% FA and 0.02% TFA. The column was operated at 45 C.

| Solvent B in % | Gradient time: |  |  |  |  |
| --- | --- | --- | --- | --- | --- |
|  | **5 min** | **15 min** | **30 min** | **45 min** | **60 min** |
| 1 | 0 min | 0 min | 0 min | 0 min | 0 min |
| 7 | 1 min | 1 min | 1 min | 1 min | 1 min |
| 45 | 6 min | 16 min | 31 min | 46 min | 61 min |
| 90 | 7 min | 17 min | 32 min | 47 min | 62 min |
| 90 | 9 min | 19 min | 34 min | 49 min | 64 min |

**Table S2.** Settings of the default proteomics method and optimized parameters for the developed glycoproteomics method.

| **Parameter** | **Proteomics method** | **Optimized glycoproteomics method** |
| --- | --- | --- |
| Collision energy as % of proteomics method | + 0% | + 50 % CE |
| Collision cell RF [Vpp] | 1500 | 1700 |
| Prepulse storage [µs] | 12 | 10 |
| Transfer time [µs] | 60 | 60 |
| Target intensity PASEF [counts] | 20.000 | 25.000 |
| TIMS isolation width [ms] | 2.75 | 7.50 |
| Polygon coordinates for precursor selection | 150 m/z @ 0.6 1/K_0_, 1300 m/z @ 1.6 1/K_0_, 1800 m/z @ 1.6 1/K_0_, 1800 m/z @ 0.6 1/K_0_ | 600 m/z @ 1.0 1/K_0_, 1900 m/z @ 1.8 1/K_0_, 2200 m/z @ 1.8 1/K_0_, 2200 m/z @ 1.3 1/K_0_, 1400m/z @ 0.7 1/K_0_, 600 m/z @ 0.7 1/K_0_ |
| Mass range [m/z] | 100-1700 | 50-4000 |
| Mobility range [1/K_0_] | 0.6-1.6 | 0.7-1.5 |
| Delta potentials [V] | D1: -20  D2: -160  D3: 110  D4: 110  D5: 0  D6: 55 | D1: -20  D2: -160  D3: 110  D4: 110  D5: 0  D6: 55 |
| Ramp time [ms] | 100 | 100 |
| Accumulation time [ms] | 100 | 100 |

**Table S3.** Optimization scheme.

| **Optimization step** | **Nanobooster fill level (ACN) [mL]** | **CE energy** | **Collision cell RF [Vpp]** | **Prepulse storage [µs]** | **Transfer time [µs]** | **Target intensity PASEF** | **Tims isolation width [ms]** | **Polygon** | **Gradient length [min]** | **Mobility range [1/K_0_]** |
| --- | --- | --- | --- | --- | --- | --- | --- | --- | --- | --- |
| CE energies | 600 | x | 1500 | 12 | 100 | 100 K | 2.75 | Default | 15 | 0.6-1.6 |
| Collision cell RF [Vpp] | 600 | +50 % | x | 12 | 80 | 100 K | 2.75 | Default | 15 | 0.6-1.6 |
| Prepulse storage [µs] | 600 | +50% | 1500 | x | 80 | 100 K | 2.75 | Default | 15 | 0.6-1.6 |
| Transfer time [µs] | 600 | +50% | 1500 | 12 | x | 100 K | 2.75 | Default | 15 | 0.6-1.6 |
| Target intensity PASEF | 600 | +50% | 1700 | 10 | 80 | x | 2.75 | Default | 15 | 0.7-1.5 |
| Tims isolation width [ms] | 600 | +50% | 1700 | 10 | 60 | 25 K | x | Default | 15 | 0.7-1.5 |
| Polygon | 600 | +50% | 1700 | 10 | 60 | 25 K | 7.5 | X | 15 | 0.7-1.5 |
| Gradient length [min] | 600 | +50% | 1700 | 10 | 60 | 25 K | 7.5 | Optimized | x | 0.7-1.5 |
| Benchmarking |  |  |  |  |  |  |  |  |  |  |
| Proteomics method (default) | No DEN (Air) | + 0 % | 1500 | 12 | 60 | 20 K | 2.75 | Default | 30 | 0.6 – 1.6 |
| Proteomics method (default) + DEN | 600 | + 0 % | 1500 | 12 | 60 | 20 K | 2.75 | Default | 30 | 0.6 – 1.6 |
| Glycoproteomics methods | No DEN (Air) | + 50 % | 1700 | 10 | 60 | 25 K | 7.5 | Optimized | 30 | 0.7 – 1.5 |
| Glycoproteomics methods + DEN | 600 | + 50 % | 1700 | 10 | 60 | 25 K | 7.5 | Optimized | 30 | 0.7 – 1.5 |
